# Nutritional ecology provides insights into competitive interactions between closely-related marten species

**DOI:** 10.1101/599084

**Authors:** Andrea Gazzola, Alessandro Balestrieri

## Abstract

For generalist, closely-related predators as those belonging to the genus *Martes*, it is a hard task to differentiate the effects on feeding habits of variation in food availability from those of resource competition. To overcome this obstacle, we reviewed dietary studies that assessed the relative bulk of each food item, as either percent biomass or percent mean volume, in the diet of both the pine-(*M. martes*) and stone-(*M. foina*) marten, and calculated the nutrient profiles (intakes of protein, lipids and carbohydrates, expressed as percentages of total metabolizable energy) of each diet. Both martens’ diets tightly clustered (average values: 47% protein-, 39% lipid- and 14% carbohydrate energy), but, most interestingly, in allopatry the nutritional niches of the two species did not differ, while the stone marten ate more carbohydrates and less protein when sympatric with the pine marten. Our data suggest that stone marten frugivory is the result of interspecific competition.

## Introduction

Pine marten *Martes martes* and stone marten *Martes foina* are the most similar European carnivores (see [1]) and co-exist over a large area, including most of central and southern Europe [2]. While forested habitats are key-features for both species, the stone marten often occurs in rural and suburban areas [3,4]. This association with human-dominated habitats has been explained as the result of competition with the pine marten [5], and may be enhanced by stone marten’s higher tolerance towards human disturbance [6]. Nonetheless, in the last two decades the pine marten has been reported to occur also in rural [7,8] and even intensively cultivated areas [9,10].

Although spatial segregation should entail the use of partially different food resources [11,12], the two martens’ trophic niches overlap extensively [13,14]. Both species are generalist feeders, using a large variety of food items, with small mammals and fruit often forming the bulk of their diets (reviewed by [15–18]). These diverse feeding habits make it hard to compare the food preferences of the two martens at range scale. Moreover, as their faeces cannot be distinguished by eye, the diets of the two martens have mostly been studied separately, preventing from differentiating the effects of local and temporal variation in food availability from those of interspecific (or intra-guild) competition [13].

Recently, Machovsky-Capuska et al. [19] suggested that a deeper insight into feeding ecology can be obtained by a multidimensional approach, integrating the percentage contribution of food items to diets with their nutritional composition. Animals typically regulate the amounts and balance of macronutrients (i.e. protein, lipids and carbohydrates) in their diets by either selecting nutritionally balanced foods or combining complementary foods to achieve a species-specific macronutrient *intake target* [20,21,22]. Previously confined to laboratory experiments with captive animals and controlled diets, the assessment of the macronutrient composition of meso-carnivore diets has been demonstrated to be achievable by the analysis of stomach contents or faecal samples from wild-living animals [23]. Till now this indirect method has been successfully applied to a handful of species, including the pine marten [24].

We hypothesized that macronutrient gains may enable us to compare the feeding requirements of the pine-and stone martens independently from the way their populations respond to various food availabilities and thus more effectively than by classical estimates of trophic niche overlap at food-level. To test for this hypotheses, we reviewed available data on the foods eaten by pine- and stone marten populations across Europe and estimated the percentage of macronutrients in each diet to quantify and compare their nutritional niches.

## Methods

We selected those studies that assessed, as either per cent biomass or per cent mean volume, the relative bulk of each food item in marten diet by the analysis of either stomach or faecal samples. Minimum sample size was set at 60 [17,24] and studies had to cover at least one complete year. When only seasonal data were reported, the mean annual relative bulk of each food item was calculated *a posteriori*. Available literature (e.g. [23,25,26]) and on-line databases (e.g. https://ndb.nal.usda.gov/; www.valori-alimenti.com) were checked to obtain, on a wet weight basis, the macronutrient composition (mean percentage of protein, lipids, and carbohydrates) of the food items used by martens (Tab. S1).

Nutrient profiles were calculated following Remonti et al. [23,24] and expressed as percentages of total metabolizable energy (kcal) using the following coefficients: protein = 14.64 kJ/g; fat = 35.56 kJ/g; non-structural carbohydrates = 14.64 kJ/g [27,28]. Macronutrient compositions of martens’ selected diets were visualized by equilateral mixture triangles (EMTs), which allow to represent 3-components (i.e. protein, lipids and carbohydrates) in a two-dimensional nutrient space [27].

To assess the effects of interspecific competition on macronutrient intakes, data were split into four groups, resulting from range overlap (sympatry *vs*. allopatry).

Finally, based on the description of study areas in selected studies, data were also split in three habitat types, broadly arranged along a gradient of increasing human interference: i) temperate forests or Mediterranean shrubland; ii) forest (shrubland) – agricultural mosaics; iii) agricultural areas.

Nutritional niche overlap between pine- and stone marten was assessed by *dynamic range boxes* (*dyn*RB; [29]), a non-parametric, robust approach for analysing compositional data [30]. Trait space overlap *port (a, b)* was calculated by the aggregation methods “product”, which calculates the geometric volume delineated by the interval lengths of each dimension of Hutchinson’s hypervolume (the sides of the boxes), and ‘‘mean’’ (of the side lengths), which assesses how similar two niches are based on the mean overlap of the *n* dimensions [29]. Data analyses were performed using R package *dyn*RB [31].

Macronutrient intakes were then compared by permutational multivariate analysis of variance (PERMANOVA; [32]) with 10000 permutations. Analyses were performed in R (3.5.1 version), with the package “vegan” [33]. Mann-Whitney (*U*) or Kruskal-Wallis tests were used as post-hoc tests to compare single nutrients, while raw frequency data of food items in the two diets were compared by the chi-squared (*χ*^2^) test.

## Results

Thirty-two studies (mean sample size: 507.7, min-max: 73-2449) were selected (Tab. S1), covering the two martens’ European ranges from Portugal in the west to Belarus in the east and from Sweden in the north to southern Spain (Fig. S1).

Mammals, particularly small rodents and insectivores, and fruit formed the bulk of both marten diets, followed by birds. The contribution of major food items to marten diets varied largely (small mammals: 7-81.2% for the stone marten and 15.5-74.8% for the pine marten; fruit: 9-63.5% and 3-54%, respectively). The total number of mammal prey recorded was 41 (stone marten: 33; pine marten: 27), with the pine marten relying on large mammals more frequently than the stone marten (*χ*^2^ = 4.41, *p* = 0.03), which, on its turn, ate more synanthropic rodents (*χ*^2^ = 5.5, *p* = 0.02). Both martens used a large variety of fruits (34 species, 29 for the stone marten and 22 for the pine marten), with the stone marten eating cultivated species more often than the pine marten (*χ*^2^ = 31.3, *p* < 0.001).

Mean (±SE) macronutrient intakes (Tab. S2; Fig. 1) were 48.9 ± 1.5% protein-, 40.3 ± 0.8%, lipid- and 10.8 ± 1.9% carbohydrate energy for the pine marten and 44.4 ± 1.7% protein-, 37.0 ± 1.3%, lipid- and 18.6 ± 2.6% carbohydrate energy for the stone marten. Lipid- and carbohydrate intakes differed significantly between species (*U* = 59.0, *p* = 0.011 for both), nonetheless the average protein:lipid ratio did not differ (1.22 ± 0.03 and 1.21 ± 0.04, respectively).

**Figure 1.**
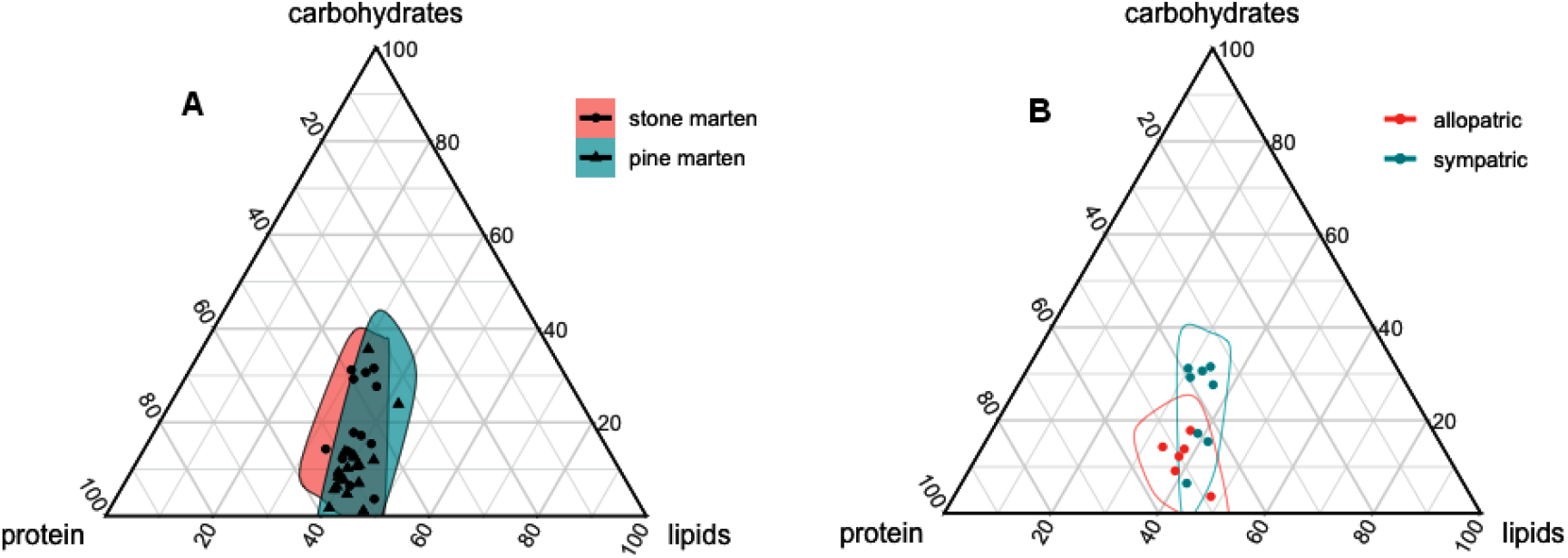
Equilateral mixture triangle showing the estimated macronutrient intake of A) pine- and stone martens from 32 European sites, and B) the stone marten living in either allopatry or sympatry with the closely-related pine marten. Data are expressed as percentage of metabolizable energy.

The largest divergence in nutritional niches occurred between allopatric and sympatric stone martens, while, on the opposite, pine marten niches overlapped almost completely (Fig. 2A). The niche of allopatric pine marten overlapped that of allopatric stone marten to a larger extent that the opposite (*a* over *b* vs *b* over *a*), consistently with allopatric stone marten showing the smallest niche size (Fig. S2). The aggregation method “mean” pointed out that the overlaps between the trait spaces of sympatric pine- and stone marten and vice-versa were intermediate and nearly symmetric (Fig. 2B). Accordingly, analysis of variance showed that pine marten macronutrient intake did not vary between allopatry and sympatry (*F* = 0.001, *p* = 0.91), while, on average, the stone marten consumed less protein and more carbohydrates in sympatry (*F* = 7.25, *p* = 0.02; Tab. 1; Fig. 1). The average macronutrient compositions of the two martens’ diets in allopatry did not differ (*F* = 0.69, *p* = 0.54; Tab. 1), while differed in sympatry (*F* = 8.17, *p* = 0.01).

**Table 1.** Average macronutrient intakes of pine- and stone martens occurring in either allopatry or sympatry.

| Species | Range | Protein | Lipids | Carbohydrates |
| --- | --- | --- | --- | --- |
| Pine marten | allopatry | 49.3 | 40.4 | 10.3 |
|  | sympatry | 48.7 | 40.2 | 11.1 |
| | | $U = 34.0, p = 0.68$ | $U = 38.0, p = 0.96$ | $U = 38.0, p = 0.96$ |
| Stone marten | allopatry | 49.4 | 38.7 | 11.8 |
|  | sympatry | 42.5 | 36.4 | 21.1 |
| | | $U = 4.0, p = 0.01$ | $U = 17.0, p = 0.37$ | $U = 7.0, p = 0.03$ |

**Figure 2.**
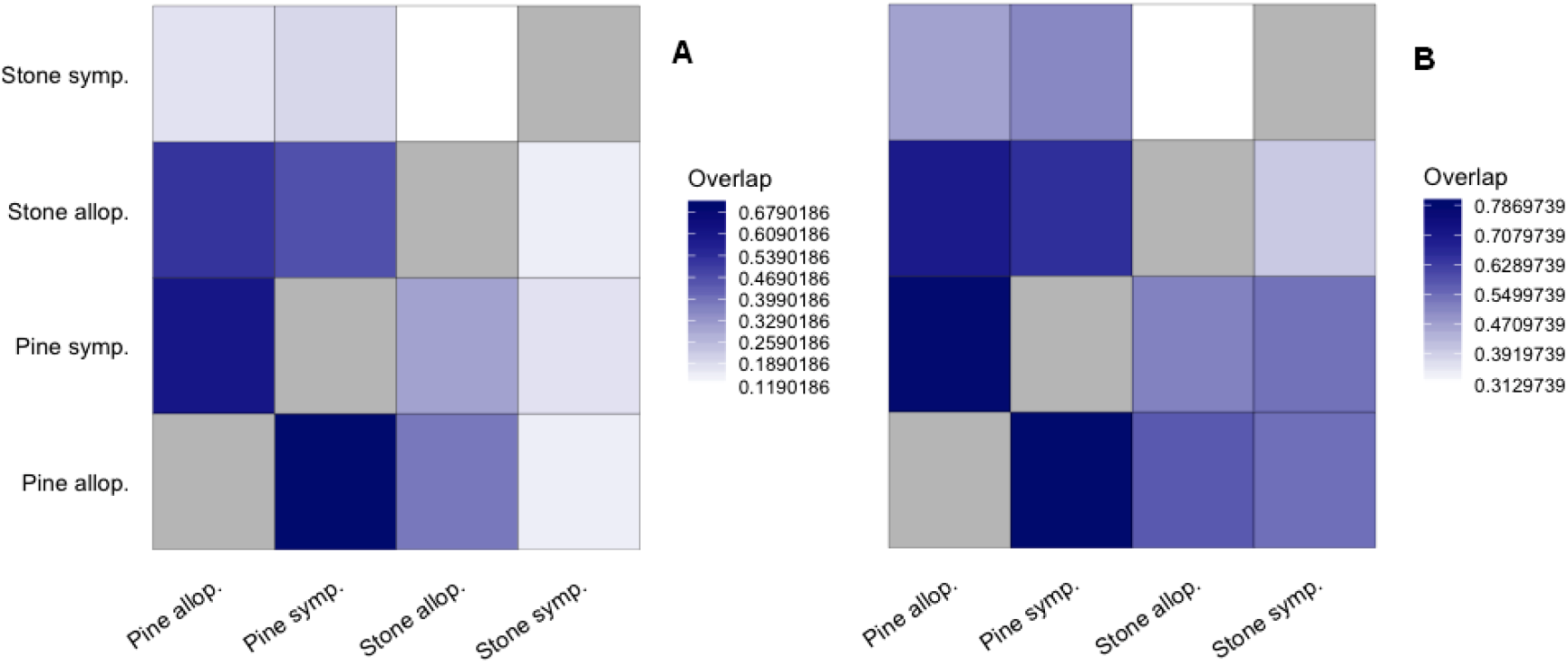
Nutritional niche overlap of pine- and stone marten in different conditions (sympatry vs. allopatry), as calculated using dynamic range boxes and the aggregation methods ‘product’ (A) and ‘mean’ (B). Heatmaps depict the overlaps *port* (*a, b*) between the nutrient spaces of species *a* and *b* based on *n* = 3 macronutrients. Light colours indicate little or no overlap.

Most studies on pine marten (77.8%) were carried out in forests, preventing from comparing diets at habitat-level, while no significant difference was recorded in the macronutrient intakes of stone martens living in different habitats (Kruskal-Wallis test, χ^2^ = 0.1-0.6; *p* > 0.7 for all tests).

## Discussion

Despite we reviewed literature studies which aimed to assess the relative importance of food items rather than macronutrient intakes, and inevitably varied in the type and quality of laboratory procedures, the diets of both martens tightly clustered. The similarity in the macronutrient composition of a large variety of diets throughout the wide European range of each species confirms that both martens tend to balance their nutrient intake by combining available food resources appropriately.

Most importantly, we demonstrated that in allopatry the nutritional niches of the two species do not differ, an achievement previously hindered by between-species differences in diet composition due to geographic variation in food availability. Our results suggest that these two mustelids, extremely similar in size, physiology and hunting strategies, also share almost identical nutrient requirements. While sympatry does not seem to involve significant deviations from the macronutrient intake target of the pine marten, we recorded an increase in the percent contribution of carbohydrates (i.e. fruit) in the diet of the stone marten, to the detriment of protein gain. Dietary shift in the stone marten may depend on two alternative mechanisms: 1) higher tolerance to carbohydrates respect to the pine marten, which would imply a larger variety of food items potentially exploitable (second and third level, respectively, of the nutritional niche size proposed by Machovsky-Capuska et al. [19]), or 2) higher competitive ability of the pine marten, which would force the stone marten to use less suitable foods.

The first hypothesis is not supported by available data: high carbohydrate levels have been reported also for the pine marten, in habitats that can be regarded as sub-optimal for this species [24,34,35]. Moreover, while the pine marten was reported to be the least frugivorous mesocarnivore of Mediterranean Europe [36], our review has pointed out that both martens can use a large variety of berries.

The competition hypothesis is consistent with the capacity of the pine marten to exclude the stone marten from forested habitats, which has been suggested by Delibes [5] as evidence of pine marten dominance. Spatial segregation *per se* can affect diet composition, by giving access to different food resources. As an example, in agricultural areas, cereals provide one-third of the protein intake of European badgers (*Meles meles*), with the consequence of an increase in carbohydrate consumption with respect to forested habitats [23]. The use of cultivated fruit and synanthropic rodents by the stone marten clearly reflects its occurrence in anthropogenic habitats (as so as pine marten’s preference for large mammals depends on its more northern range; [16]). Nonetheless, the average macronutrient balance of the stone marten did not vary among habitats, suggesting that diet shift may also result from fine-grained adjustments in the use of space (e.g. use of less suitable, in terms of food availability, but pine marten-free food patches; see [37]) or differential use of food resources in overlapping hunting areas [38]. Accordingly, in allopatry the protein intake of the stone marten was high (45-52%) in very different habitats, from intensively cultivated lowland to Alpine forests and Mediterranean scrubland. Interestingly, at the southern edge of pine marten range, the pattern of dominance is reversed and the pine marten is more frugivorous than the more abundant and spread stone marten [35], supporting the role played by interspecific competition in shaping the diet of these mustelids. However, *dyn*RB aggregation method “mean” suggests that diet shifts do not differentiate the nutritional niches of sympatric marten species to the extent necessary to prevent competition, implying that co-existence is the results of a mosaic of behavioural adjustments, including habitat-[4] and time-partitioning [39].

For the first time, nutritional ecology provided evidence that stone marten frugivory may be, at least partially, a makeshift strategy to cope with competition with the pine marten, rather than the result of the selection for a highly profitable resource. To paraphrase Rosenzweig [40], from a macronutrient perspective the stone marten would do better, on average, investing its time and energy to pursue protein-rich small vertebrates than searching for fruit, but this alternative food resource may enhance its coexistence with the pine marten throughout Europe. We argue that our approach may contribute to understand the feeding strategies of other closely-related sympatric species, provided that dietary studies are fostered and data expressed in terms of volume or biomass.

---

**Figure S1.**
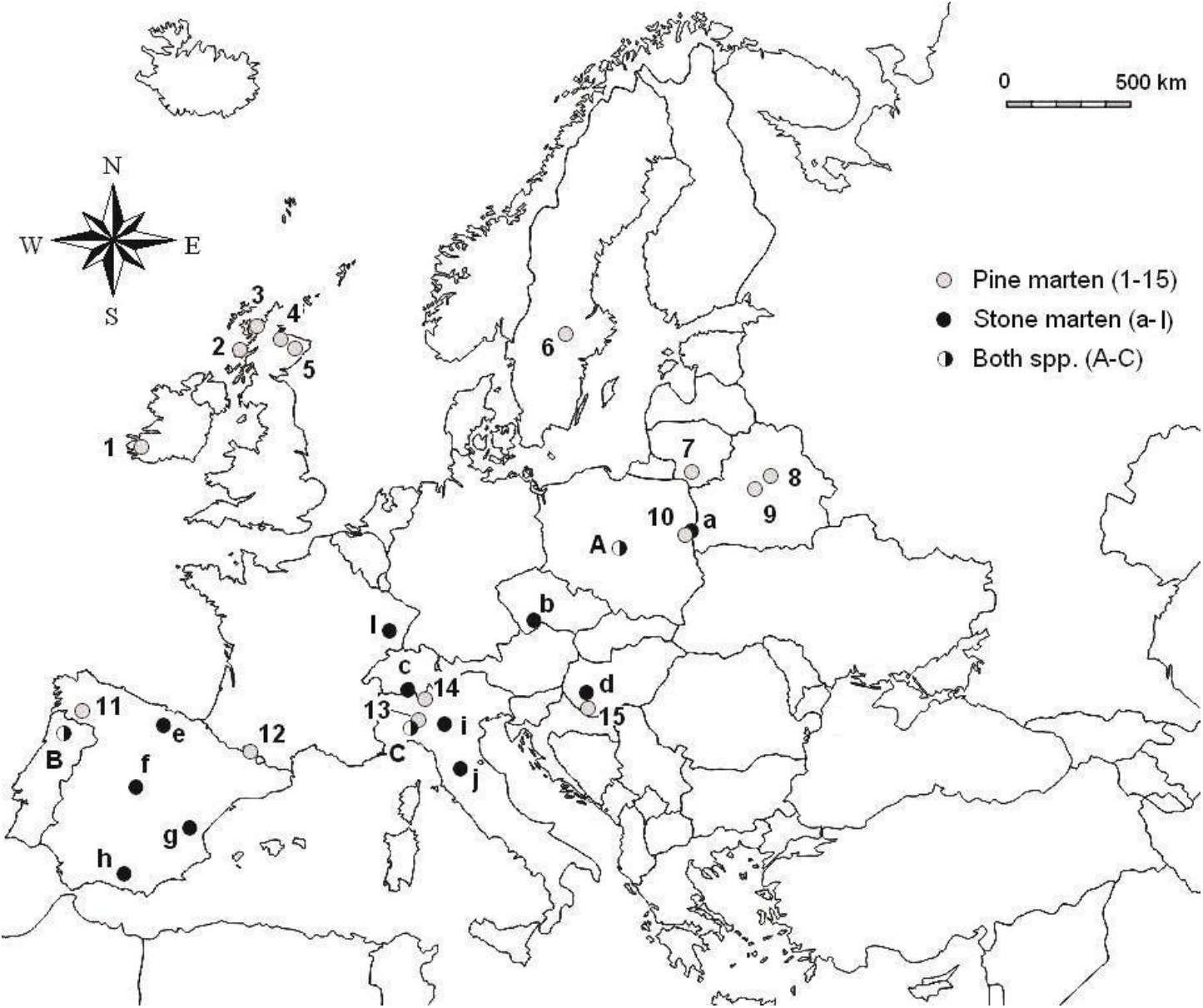
Distribution of the 32 reviewed study areas in Europe (letters and numbers correspond to those in Tab. 1).

**Figure S2.**
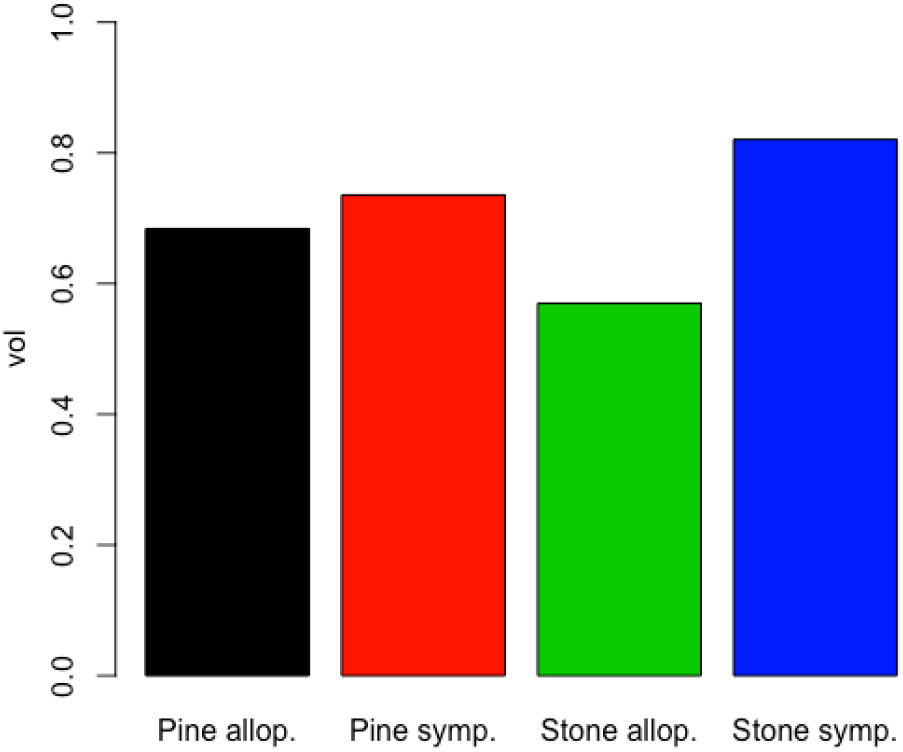
Sizes (*vol*) of nutritional spaces of pine- and stone marten in different conditions (sympatry vs. allopatry).

**Table S1.** Macronutrient content of martens’ food resources.

| Food items | Protein | Lipids | Carbohydrates |
| --- | --- | --- | --- |
| **Fungi** | 3.9 | 0.7 | 0 |
| **Fruit** | 1.1 | 0.9 | 18.1 |
| Wild Rosaceae | 1.6 | 0.5 | 18.5 |
| *Rubus* spp. | 1.3 | 0 | 5.7 |
| *Rosa* spp. | 1.6 | 0.3 | 38.2 |
| *Prunus* spp. | 0.5 | 0.1 | 10.5 |
| *Sorbus aucuparia* | 2.3 | 2.0 | 14.1 |
| *Crataegus monogyna* | 2.5 | 0.0 | 24.2 |
| Cultivated Rosaceae | 0.4 | 0.3 | 10.0 |
| *Mespilus germanica* | 0.4 | 0.4 | 6.1 |
| *Prunus domestica* | 0.7 | 0.3 | 11.4 |
| *Malus communis* | 0.3 | 0.1 | 13.7 |
| *Pirus communis* | 0.3 | 0.4 | 8.9 |
| *Diospyros kaki* | 0.6 | 0.3 | 17.0 |
| *Hedera ilex* | 1.8 | 5.1 | 21.0 |
| *Arbutus unedo* | 1.0 | 0.0 | 11.0 |
| *Vaccinium* spp. | 0.9 | 0.2 | 5.1 |
| *Ribes* spp. | 0.9 | 0.0 | 6.6 |
| *Vitis vinifera* | 0.5 | 0.1 | 13.5 |
| *Ficus carica* | 0.8 | 0.3 | 19.2 |
| *Taxus baccata* | 2.1 | 3.9 | 5.6 |
| *Castanea sativa* | 1.8 | 2.9 | 39.4 |
| *Juniperus communis* | 0.5 | 0.5 | 80.0 |
| Earthworms | 11.8 | 1.5 | 0.0 |
| **Insects** | 20.3 | 8.6 | 0.0 |
| Larvae | 17.8 | 13.3 | 0.0 |
| **Reptiles** | 19.8 | 3.7 | 0.0 |
| **Amphibians** | 18.3 | 3.7 | 0.0 |
| **Birds** | 21.7 | 5.4 | 0.1 |
| **Mammals** | 19.7 | 8.5 | 0.0 |
| *Lepus europaeus* | 21.8 | 2.3 | 0.0 |
| *Oryctolagus cuniculus* | 20.0 | 4.0 | 0.0 |
| Insectivores | 19.2 | 5.9 | 0.0 |
| Rodents | 19.8 | 9.0 | 0.0 |
| *Hystrix/Myocastor* | 20.1 | 10.2 | 0.0 |
| Small rodents | 19.6 | 8.1 | 0.0 |
| *Rattus* sp. | 20.1 | 10.2 | 0.0 |
| Wild ungulates | 22.3 | 2.9 | 0.0 |
| Livestock (beef and sheep) | 19.0 | 15.0 | 0.0 |
| Carnivores | 15.5 | 16.9 | 0.0 |
| Eggs | 13.0 | 11.0 | 0.4 |
| Trout *Salmo trutta* | 1.7 | 14.7 | 3.0 |
| Garbage | 10.5 | 3.1 | 8.8 |

**Table S2.** Macronutrient (protein, lipids and carbohydrates) intakes of pine- (PM) and stone martens (SM), as assessed for the 32 selected studies. Data are expressed as percentage of metabolizable energy (N: sample size; codes correspond to those in Figure S1).

| Spp. | N | Protein | Lipids | Carboh. | code | Range | Habitat | References |
| --- | --- | --- | --- | --- | --- | --- | --- | --- |
| PM | 335 | 52.8 | 38.9 | 8.3 | 7 | allopatry | mosaic | Baltrunaite 2002 |
|  | 2449 | 47.7 | 41.5 | 10.8 | 4 |  | forest | Caryl et al. 2012 |
|  | 760 | 52.1 | 40.1 | 7.7 | 6 |  | forest | Helldin 2000 |
|  | 546 | 57.8 | 40.5 | 1.7 | 5 |  | forest | Kubasiewicz 2014 |
|  | 337 | 52.3 | 38.4 | 9.3 | 3 |  | forest | Lockie 1960 |
|  | 387 | 33.9 | 42.2 | 23.9 | 1 |  | forest | Lynch & McCann 2007 |
|  | 164 | 48.3 | 41.1 | 10.5 | 2 |  | forest | Putman 2000 |
|  | 109 | 48.9 | 37.4 | 13.7 | 13 | sympatry | agriculture | Balestrieri et al. 2011 |
|  | 148 | 47.7 | 39.3 | 12.9 | 14 |  | forest | Biancardi & Rinetti 2002 |
|  | 1735 | 51.8 | 47.1 | 1.1 | 10 |  | forest | Jedrzejewski et al. 1993 |
|  | 332 | 53.0 | 42.3 | 4.7 | 15 |  | forest | Lanszki et al. 2007 |
|  | 97 | 33.6 | 30.8 | 35.6 | B |  | forest | Monterroso et al. 2016 |
|  | 155 | 49.6 | 43.3 | 7.1 | A |  | mosaic | Posluzny et al. 2007 |
|  | 109 | 50.1 | 39.6 | 10.3 | C |  | agriculture | Remonti et al. 2012 |
|  | 198 | 47.7 | 39.7 | 12.6 | 11 |  | forest | Rosellini et al. 2007 |
|  | 445 | 44.4 | 43.6 | 11.9 | 12 |  | forest | Ruiz-Olmo & Lopez-Martin 1996 |
|  | 1222 | 54.9 | 39.4 | 5.7 | 8 |  | forest | Sidorovich et al. 2005 |
|  | 1624 | 54.1 | 39.6 | 6.3 | 9 |  | forest | Sidorovich et al. 2010 |
| SM | 181 | 50.1 | 37.6 | 12.2 | i | allopatry | agriculture | Balestrieri et al. 2013 |
|  | 194 | 48.5 | 47.9 | 3.6 | c |  | forest | Balestrieri et al. 2018 |
|  | 140 | 52.4 | 38.5 | 9.1 | f |  | forest | Barrientos & Virgos 2006 |
|  | 157 | 48.3 | 37.8 | 13.8 | e |  | mosaic | Delibes 1978 |
|  | 856 | 45.2 | 37.0 | 17.8 | h |  | forest | Padial et al. 2002 |
|  | 137 | 52.1 | 33.6 | 14.3 | g |  | forest | Such & Calabuig 2002 |
|  | 984 | 38.9 | 29.9 | 31.2 | a | sympatry | forest | Czernick et al. 2016 |
|  | 320 | 44.1 | 38.7 | 17.2 | j |  | forest | Genovesi et al. 1996 |
|  | 1227 | 34.6 | 33.9 | 31.5 | d |  | agriculture | Lanszki et al. 2009 |
|  | 73 | 36.1 | 36.3 | 27.7 | B |  | forest | Monterroso et al. 2016 |
|  | 287 | 36.6 | 32.8 | 30.6 | A |  | mosaic | Posluzny et al. 2007 |
|  | 106 | 39.5 | 31.2 | 29.2 | C |  | agriculture | Remonti et al. 2012 |
|  | 108 | 51.6 | 42.0 | 6.5 | b |  | agriculture | Rysava-Novakova & Koubek 2009 |
|  | 326 | 43.2 | 41.4 | 15.4 | l |  | mosaic | Waetcher 1975 |

